# Ipsilesional motor cortex activation with high-force unimanual handgrip contractions of the less-affected limb in participants with stroke

**DOI:** 10.1101/2021.06.25.449456

**Authors:** Justin W. Andrushko, Layla Gould, Doug W. Renshaw, Shannon Forrester, Michael E. Kelly, Gary Linassi, Marla Mickleborough, Alison Oates, Gary Hunter, Ron Borowsky, Jonathan P. Farthing

**Affiliations:** College of Kinesiology, University of Saskatchewan, Saskatchewan, Canada; Department of Surgery, Division of Neurosurgery, College of Medicine, University of Saskatchewan, Saskatchewan, Canada; Department of Physical Medicine and Rehabilitation, College of Medicine, University of Saskatchewan, Saskatchewan, Canada; Department of Psychology, College of Arts and Science, University of Saskatchewan, Saskatchewan, Canada; Department of Medicine, Division of Neurology, College of Medicine, University of Saskatchewan, Saskatchewan, Canada

**Keywords:** Functional connectivity, Brain activation, Stroke, Ipsilateral hemisphere

## Abstract

Stroke is a leading cause of severe disability that often presents with unilateral motor impairment. Conventional rehabilitation approaches focus on motor practice of the affected limb and aim to suppress brain activity in the contralesional hemisphere to facilitate ipsilesional hemispheric neuroplasticity subserving motor recovery. Previous research has also demonstrated that exercise of the less-affected limb can promote motor recovery of the affected limb through the interlimb transfer of the trained motor task, termed cross-education. One of the leading theories for cross-education proposes that the interlimb transfer manifests from ipsilateral cortical activity during unimanual motor tasks, and that this ipsilateral cortical activity results in motor related neuroplasticity giving rise to contralateral improvements in motor performance. Conversely, exercise of the less-affected limb promotes contralesional brain activity which is typically viewed as contraindicated in stroke recovery due to the interhemispheric inhibitory influence onto the ipsilesional hemisphere. High-force unimanual handgrip contractions are known to increase ipsilateral brain activation in control participants, but it remains to be determined if this would be observed in participants with stroke. Therefore, this study aimed to determine how parametric increases in handgrip force during repeated contractions with the less-affected limb impacts brain activity bilaterally in participants with stroke and in a cohort of neurologically intact controls. In this study, higher force contractions were found to increase brain activation in the ipsilesional/ipsilateral hemisphere in both groups (*p* = .002), but no between group differences were observed. These data suggest that high-force exercise with the less-affected limb may promote ipsilesional cortical plasticity to promote motor recovery of the affected-limb in participants with stroke.

## Introduction

Ischemic stroke, characterized by neuronal cell death due to cerebrovascular disruptions (Yu et al., 2016), is the second leading cause of death globally and one of the leading causes of severe disability (Katan and Luft, 2018). Approximately 15 million people experience stroke annually, of which five million suffer from permanent disability (Mittmann et al., 2012). When the occluded vessel serves sensorimotor-relevant cortical or subcortical areas, sensorimotor impairments often manifest. Most often the impairment presents as an asymmetry in motor output that is lateralized to the limb(s) contralateral to the cerebral hemisphere that suffered the lesion. Identifying rehabilitation methods that have the potential to improve motor functional outcomes is of clear importance and a persistent research question.

In neurologically intact humans, voluntary movements are governed typically by contralateral hemispheric control (Borowsky et al., 2002; Cincotta and Ziemann, 2008), which is thought to have an active inhibitory effect on the ipsilateral hemisphere through transcallosal projections intended to suppress unwanted movements (interhemispheric inhibition [IHI]; (Hübers et al., 2008; Sehm et al., 2016)). However, in participants with stroke this transcallosal inhibitory control is altered and the contralesional hemisphere (opposite side to the lesion) exhibits an increased level of activation during movements with the affected limb. There is debate regarding brain stimulation and motor training paradigms that modulate either ipsilesional (same side as the lesion) or contralesional hemispheric brain activity (Buetefisch, 2015). The *interhemispheric competition model* (Kinsbourne, 1974; Murase et al., 2004; Bütefisch et al., 2008; Grefkes et al., 2008; Nowak et al., 2009) suggests bi-directional changes to IHI post-stroke, whereby the ipsilesional hemisphere exhibits a reduced capacity to inhibit brain activity in the non-affected contralesional hemisphere. In contrast, when the contralesional hemisphere is active it appears to have an increased capacity to inhibit the damaged hemisphere (Kinsbourne, 1974; Murase et al., 2004; Bütefisch et al., 2008; Grefkes et al., 2008; Nowak et al., 2009). Due to the changes in inhibitory control, this model suggests contralesional brain activity is maladaptive and ‘competes' with the ipsilesional hemisphere for cortical control over movement. Therefore, the interhemispheric competition model suggests that contralesional brain activity reduces the opportunity for ipsilesional neuroplasticity that would promote contralateral motor control as is seen in neurologically intact individuals. Based on this model, conventional treatments for improving motor impairment focus on finding ways to inhibit contralesional and promote ipsilesional brain activity that would then subserve preferential neuroplasticity and recovery of the ipsilesional hemisphere. However, the premise based on the interhemispheric competition model that all therapies should focus on inhibiting contralesional brain activity is somewhat controversial, as good functional motor recovery can occur in individuals that display sustained contralesional brain activation during affected limb movements (see review by (Dodd et al., 2017)).

There is evidence supporting the implementation of contralesional, less affected limb approaches in participants with chronic stroke to improve motor recovery (Dragert and Zehr, 2013; Urbin et al., 2015; Sun et al., 2018; Dehno et al., 2021). These studies are based on the concept of cross-education - a neuromuscular phenomenon referring to the increased motor output (i.e., force generation or skill-based movements) of the opposite, untrained limb following a period of unilateral motor training (Manca et al., 2021). The first empirical evidence of cross-education dates back to a case study in 1894 (Scripture et al., 1894), yet the neural mechanisms driving the effect are not completely understood. The implementation of cross-education into the rehabilitation process for promoting recovery of the more-affected limb after stroke paradoxically conflicts with the interhemispheric competition model. Given that voluntary exercise with the less-affected limb promotes brain activity in the contralesional hemisphere, it presumably would inhibit ipsilesional brain plasticity. Yet, the exemplar studies cited above suggest motor recovery is enhanced.

A leading theory describing the neural underpinnings of cross-education is the cross-activation hypothesis (Lee and Carroll, 2007; Ruddy and Carson, 2013). The cross-activation hypothesis suggests unilateral voluntary movements facilitate bilateral brain activation, which results in motor training-related neuroplasticity in both hemispheres (Lee et al., 2010; Ruddy and Carson, 2013; Manca et al., 2018). In neurologically intact humans, previous research has observed ipsilateral brain activation with high-force (Andrushko et al., 2021), fatiguing (Benwell et al., 2006; Jiang et al., 2012), or complex (Verstynen et al., 2005) unilateral voluntary movements. Further, greater cross-education is known to occur when the exercise is performed at high intensities (Urbin et al., 2015), high velocities (Farthing and Chilibeck, 2003), and when eccentric muscle actions are incorporated into the unilateral training regimen (Farthing and Chilibeck, 2003; Manca et al., 2017). Therefore, if unilateral motor tasks with the less-affected limb result in ipsilesional brain activation in individuals with stroke, targeted exercise with the less-affected limb provides a potential avenue for enhancing neuroplasticity subserving motor recovery, while shedding light on the efficacy of cross-education as an adjunct therapy for individuals with stroke (i.e., in addition to constraint-induced-movement-therapy).

The purpose of this study is to determine if high-force unilateral handgrip contractions performed with the less-affected limb in participants with stroke result in increased cortical activation and functional connectivity between the primary motor cortex (M1) and the supplementary motor area (SMA) within the ipsilesional hemisphere. Further, utilizing previously published data from neurologically intact controls (Andrushko et al., 2021), a secondary purpose is to determine if unilateral handgrip contractions performed with the less-affected limb result in differences in ipsilateral/ipsilesional brain activation and functional connectivity between participants with and without a history of stroke. The primary hypotheses are that high-force unilateral handgrip contractions will result in greater brain activation and functional connectivity within the ipsilesional hemisphere in both groups.

## Methods

### Participants

Using a partial eta squared (η_p_^2^) effect size of η_p_^2^ = 0.439 based on a main effect of condition for the modulation of ipsilateral M1 (iM1) cortical activation between 25%, 50%, and 75% maximum voluntary contraction (MVC) from previously published work (Andrushko et al., 2021), it was determined that an estimated total sample size of eight was needed (G*Power 3.1.9.2; 1-β = 0.95, α = 0.05). Eleven participants with stroke volunteered to participate in a larger intervention (clinicaltrials.gov: NCT02948725), of which their baseline data was used in the present study (Data are presented as mean ± standard deviation; 3 female, 8 male, 10 right-handed, 1 left-handed, age: 60 ± 11 yrs, height: 173.3 ± 6.4 cm, mass: 81.7 ± 18.3 kg). Additionally, 13 neurologically intact controls (11 right-handed, 2 left-handed, age: 28 ± 6 yrs, height: 170.9 ± 9.8 cm, mass: 75.1 ± 16.7 kg) from a previously published study were used to conduct between group comparisons (Andrushko et al., 2021).

Participants with stroke were eligible to participate if they met the following inclusion criteria: 18 years or older, within 18 months of stroke recovery, medically stable, and have moderate to severe upper limb hemiparesis as diagnosed using the Chedoke McMaster Stroke Assessment. Participants with stroke were excluded if they had significant cognitive impairment or aphasia affecting understanding as assessed by a clinician, severe upper limb spasticity preventing any movement of the proximal arm and shoulder, had a diagnosis of hemorrhagic or bilateral stroke, had a history of other severe upper limb musculoskeletal injury or other neurological diseases, had intracranial metal clips or cardiac pacemaker, or anything that would preclude an MRI, and finally if participants had any condition that would preclude the participant's ability to attend follow-up visits in the opinion of the investigator.

Written informed consent was obtained from all participants prior to participating. This study conforms to the standards set out by the Declaration of Helsinki and was approved by the University of Saskatchewan Biomedical Research Ethics Board (Ethics # Bio 16-157).

### Functional assessment

The participants with stroke completed the Fugl Meyer upper limb assessment (Sullivan et al., 2011) and the Chedoke-McMaster impairment inventory: stage of recovery for arm and hand (Gowland et al., 1993) to determine motor function in the more-affected limb. Additionally, the Waterloo handedness questionnaire was used to acquire an estimate of their hand dominance pre-stroke (Bryden, 1977).

### Experimental outline

Participants attended one fMRI session where they completed three experimental conditions in a randomized order, each condition involved repeated submaximal unimanual isometric handgrip contractions at either 25%, 50%, or 75% of MVC with their less-affected hand (stroke participants) or their right hand for the neurologically intact control participants. The session began with a structural MRI brain scan, followed by three MVCs with the less-affected limb (stroke) or the right hand of the control participants. One minute of rest was given between each MVC attempt to avoid fatigue. The MVCs were used to set the target force for each of the three submaximal conditions. Next, participants performed the three experimental conditions in a randomized order with a functional brain scan occurring during the performance of each condition.

### Behavioural motor task

Participants performed 5 sets × 5 repetitions of grip contractions at each prescribed submaximal force during separate scanning runs with an MRI-compatible hand clench dynamometer (Biopac Systems Inc. Aero Camino Goleta, CA). In a block design, task blocks composed of 1650 ms (i.e., corresponding to the repetition time [TR] of the T2* fMRI scan) contractions alternating with 1650 ms of rest (16500 ms total task block), separated by rest blocks of complete rest (16500 ms total rest block). During scans the participants wore MRI compatible goggles and viewed a projection of a computer screen running a custom-built LabView (version 8.6) interface. Participants saw clear target lines and go/no-go flashing lights and were cued when to contract or relax. The LabView interface was triggered by the MRI to ensure the task was synchronized with each TR. Target lines were presented relative to the individual's peak MVC and force feedback was presented as a vertical force bar that was responsive to each participant's grip contraction (i.e., harder contraction resulted in the bar rising vertically). Two virtual ‘lights' were present on the motor task interface to cue participants. A light turned green to instruct the participant to contract to the target line and turned black to indicate when to stop contracting. A second light remained black during task blocks and turned red during rest blocks to indicate a sustained rest. The red light switched to black moments before the next task block as an indicator that the next task series of contractions was about to begin. During each contraction force condition, participants were instructed to relax their non-active more-affected arm and hand to prevent mirror activity. Using this same experimental paradigm, the muscle activity of non-active arm was not significantly active compared to baseline noise in an electromyography control experiment with young neurologically intact adults (Andrushko et al., 2021).

### Handgrip motor task processing

The handgrip force data was processed in Matlab using custom scripts (The Mathworks Inc, 2018). Prior to analysis, data were processed with a fourth-order 100 Hz low pass Butterworth filter and full-wave rectified. Next the onset and offset of each contraction was determined and the mean of each contraction was calculated. Data were normalized to the mean Kg-force of the highest MVC and expressed as a percentage of MVC. The mean of each task block (five contractions) was calculated and used in subsequent analyses.

### fMRI Parameters

All scans were done in a Siemens 3T MAGNETOM Skyra MRI scanner (Siemens Healthcare, Erlangen, Germany). At the start of the session the whole-brain anatomical scan was acquired using a high-resolution magnetization prepared rapid acquisition gradient echo (MPRAGE) sequence consisting of 192 T1-weighted echo-planar imaging slices (1 mm slice thickness with no gap), with an in-plane resolution of 1 × 1 mm (field of view = 256 × 256; TR = 1900 ms; echo time [TE] = 2.08 ms). For each of the experimental conditions T2*-weighted single-shot gradient-echo planar imaging scans were acquired using an interleaved ascending sequence, consisting of 105 volumes (TR = 1650 ms; TE = 30 ms) of 25 axial slices of 4-mm thickness (1-mm gap) with an in-plane resolution of 2.7 mm × 2.7 mm (field of view = 250) using a flip angle of 90°. The top 2 coil sets (16 channels) of a 20-channel Siemens head-coil (Siemens Healthcare) were used. Scans consisted of a 10-volume alternating block design beginning with five volumes for stabilization (task, rest; 105 volumes total).

### fMRI Pre-Processing

Functional MRI data processing was carried out using FMRI Expert Analysis Tool (FEAT) Version 6.00, as part of FSL (FMRIB's Software Library, www.fmrib.ox.ac.uk/fsl). Boundary based registration was used to register the functional image to the high-resolution T1-weighted structural image. Registration of the functional images to the T1-weighted structural image was carried out using FLIRT: (Jenkinson and Smith, 2001; Jenkinson et al., 2002), and the registration to the standard space images was carried out using FMRIB's Nonlinear Image Registration Tool (FNIRT; (Andersson et al., 2007a, 2007b)).

The following pre-processing was applied: motion correction using Motion Correction FMRIB's Linear Image Registration Tool (MCFLIRT; (Jenkinson et al., 2002)); non-brain removal using Brain Extraction Tool (BET; (Smith, 2002)); spatial smoothing using a Gaussian kernel of FWHM 6mm; grand-mean intensity normalization of the entire 4D dataset by a single multiplicative factor (Pruim et al., 2015).

Next, Independent Component Analysis Automatic Removal of Motion Artifacts (ICA-AROMA) was used to identify and remove motion-related noise from the functional data (Pruim et al., 2015). Following the ICA-AROMA data clean up, data were high pass temporal filtered with a 0.01 Hz cut off frequency. Time-series statistical analyses were carried out using FMRIB's Improved *Linear Model* (FILM) with local autocorrelation correction (Woolrich et al., 2001). Z (Gaussianised T/F) statistic images were constructed non-parametrically using Gaussian Random Field theory-based maximum height thresholding with a corrected significance threshold of *p* = 0.05 (Worsley, 2001).

#### On-task functional connectivity

To assess the on-task functional connectivity during the three different conditions, the 50 volumes corresponding with the task blocks were extracted and merged across time. Rest volumes were removed to avoid the potential impact that rest-related activity may have on the functional connectivity analysis (Steel et al., 2016; Cole et al., 2018) and the same pre-processing steps previously indicated were used on the 50 volume on-task scans. Following data processing, the mean timeseries of the ipsilesional (stroke)/ipsilateral (controls) and contralesional (stroke)/contralateral (controls) M1 and SMA (iM1, cM1, iSMA, cSMA respectively) were extracted from native-space using reverse transformed region of interest (ROI) masks from the Brainnetome atlas (left M1: A4ul_l; right M1: A4ul_r; left SMA: A6m_l; right SMA: A6m_r) in MNI standard space (Fan et al., 2016). Correlation analyses were then carried out using custom Matlab scripts (The Mathworks Inc, 2018). First a correlation matrix was calculated that represented the edge strength between each network node. Next, a Fishers r-to-z transformation was carried out on the Pearson's r values. The Z-scores for the edge strengths were then used for analyses.

### Statistical analyses

Analyses were carried out in R (R Core Team, 2019), using linear mixed effects (LME) analyses, with participants treated as random effects to account for repeated measures. The following R packages were used; lmerTest package (Kuznetsova et al., 2017), tidystats (Sleegers, 2020), ggplot2 (Wickham, 2016).

#### Handgrip force

To assess the relative handgrip force between groups and conditions, fixed effects of group (stroke, control), condition (25%, 50%, 75% MVC), and block (5 blocks of contractions during each condition), in addition to interactions of group × condition, condition × block, group × block, and a group × condition × block were included in the model.

#### Brain activation

To assess the brain activation in each hemisphere during the three different conditions for each group, two separate LME analyses were carried out for each hemisphere for the percent signal change of the M1 and SMA, with fixed effects of group, condition, and a group × condition interaction.

#### On-task functional connectivity

To assess the functional connectivity from the 50-volume ‘on-task’ data, separate analyses were carried out to examine interhemispheric homologous functional connectivity of M1's bilaterally (cM1-iM1) and SMA's bilaterally (cSMA-iSMA). Further, intrahemispheric connectivity was assessed between the M1 and SMA (M1-SMA) within each hemisphere, respectively. For each of these dependent variables, LME analyses were carried out, with fixed effects of group, condition, and a group × condition interaction included in each model.

## Results

### Motion and data removal

Data from three participants with stroke during the 75% MVC condition was deemed unusable due to high levels of motion artifact that was task correlated (table 2). An additional scan in the 25% condition was missing due to a technical error during data collection. Furthermore, of the usable scans, several data points for each ROI were not included in analyses due to lesion intrusion within the ROI masks in participants with stroke. This resulted in a total data loss of three ROIs for the iM1 (two left, one right), and two ROIs for the iSMA (two right). After data removal for the 25% MVC condition, a total of seven ROIs for the iM1, 10 for the unaffected M1, eight ROIs for the affected SMA, and 10 for the unaffected SMA were deemed usable. For the 50% MVC condition, a total of eight ROIs for the affected M1, 11 for the unaffected M1, nine for the affected SMA and 11 for the unaffected SMA were included in analyses. Finally, for the 75% MVC condition there were five usable ROIs for the affected M1, eight ROIs for the unaffected M1, six ROIs for the affected SMA, and nine ROIs for the unaffected SMA.

**Table 1.**
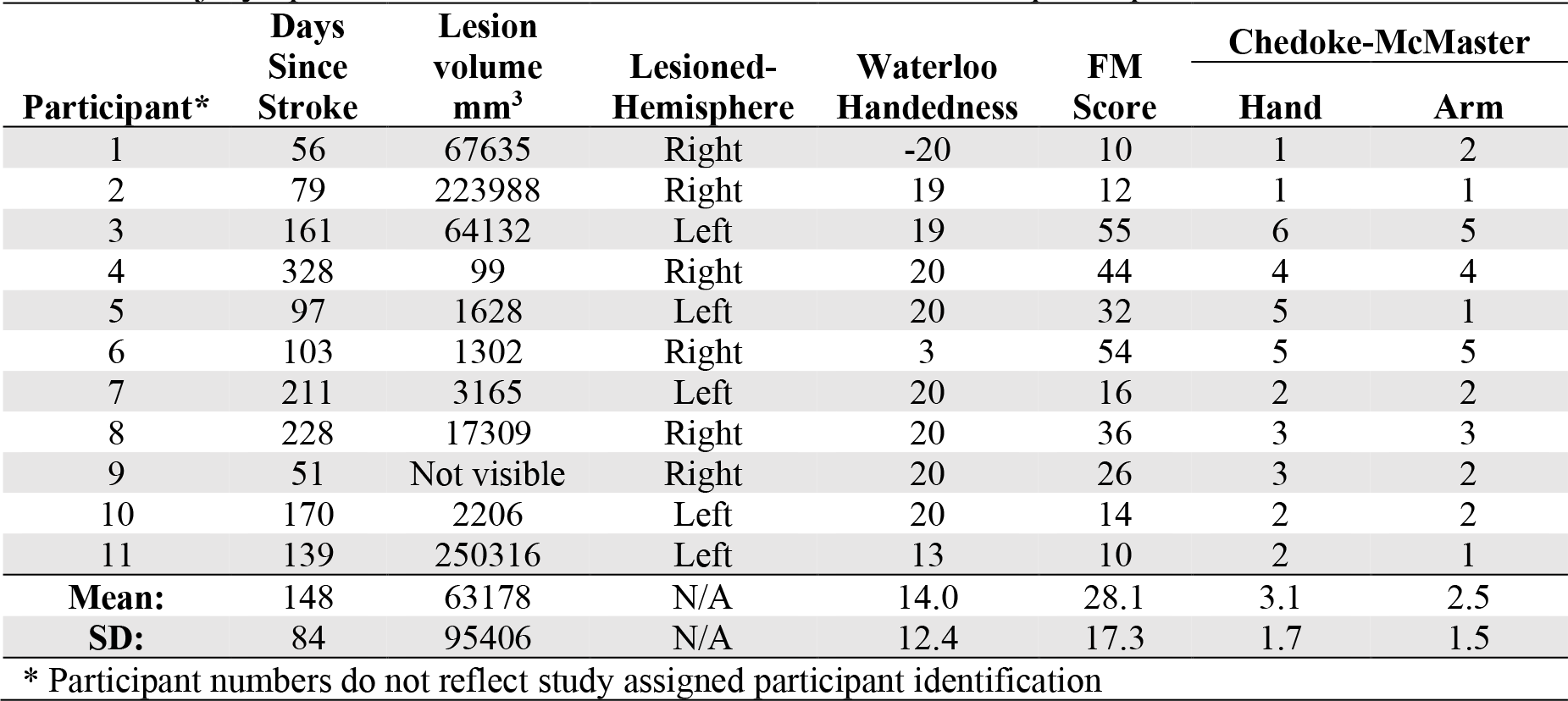
Injury-specific characteristics and functional scores for participants with stroke

**Table 2.** Motion metrics and data removal

| Participant** | Group | 25% MVC |  | 50% MVC |  | 75% MVC |  |
| --- | --- | --- | --- | --- | --- | --- | --- |
|  |  | Motion (mm) |  | Motion (mm) |  | Motion (mm) |  |
|  |  | Relative | Absolute | Relative | Absolute | Relative | Absolute |
| 1 | Stroke | 0.11 | 0.19 | 0.14 | 0.25 | 0.13 | 0.30 |
| 2 | Stroke | 0.16 | 0.15 | 0.16 | 0.14 | 0.21 | 0.44 |
| 3 | Stroke | No Scan | No Scan | 0.52 | 0.18 | 8.95* | 1.34* |
| 4 | Stroke | 0.08 | 0.12 | 0.10 | 0.45 | 0.13 | 0.29 |
| 5 | Stroke | 0.09 | 0.23 | 0.26 | 1.01 | 0.46* | 1.12* |
| 6 | Stroke | 0.05 | 0.08 | 0.11 | 0.23 | 0.29 | 0.64 |
| 7 | Stroke | 0.13 | 0.14 | 0.15 | 0.16 | 0.18 | 0.56 |
| 8 | Stroke | 0.19 | 0.35 | 0.19 | 0.27 | 0.20 | 0.22 |
| 9 | Stroke | 0.17 | 0.25 | 0.15 | 0.26 | 0.78* | 2.11* |
| 10 | Stroke | 0.16 | 0.13 | 0.15 | 0.17 | 0.19 | 0.24 |
| 11 | Stroke | 0.20 | 0.30 | 0.23 | 0.31 | 0.27 | 0.46 |
| 12 | Control | 0.05 | 0.08 | 0.07 | 0.22 | 0.10 | 0.35 |
| 13 | Control | 0.06 | 0.12 | 0.08 | 0.25 | 0.13 | 0.58 |
| 14 | Control | 0.10 | 0.28 | 0.11 | 0.36 | 0.10 | 0.25 |
| 15 | Control | 0.11 | 0.27 | 0.05 | 0.10 | 0.06 | 0.07 |
| 16 | Control | 0.03 | 0.06 | 0.04 | 0.13 | 0.06 | 0.20 |
| 17 | Control | 0.07 | 0.11 | 0.06 | 0.10 | 0.07 | 0.13 |
| 18 | Control | 0.06 | 0.13 | 0.11 | 0.31 | 0.11 | 0.22 |
| 19 | Control | 0.10 | 0.26 | 0.12 | 0.21 | 0.19 | 0.56 |
| 20 | Control | 0.05 | 0.11 | 0.06 | 0.27 | 0.11 | 0.29 |
| 21 | Control | 0.04 | 0.13 | 0.06 | 0.17 | 0.17 | 0.59 |
| 22 | Control | 0.04 | 0.11 | 0.06 | 0.21 | 0.15 | 0.73 |
| 23 | Control | 0.04 | 0.06 | 0.07 | 0.15 | 0.15 | 0.36 |
| 24 | Control | 0.04 | 0.07 | 0.05 | 0.09 | 0.06 | 0.17 |

\* Scan removed due to high motion or task correlated motion
\*\* Participant numbers do not reflect study assigned participant identification

For the relative handgrip force, a significant main effect of condition was found (*F*(2, 322) = 4012.53, *p* < .001). However, the main effects of group (*F*(1, 23) = 0.65, *p* = .43), and block (*F*(4, 322) = 0.18, *p* = .95) were not significant, indicating that relative handgrip force differed between conditions but not between groups or across the five blocks of contractions. Additionally, the group × condition (*F*(2, 322) = 1.01, *p* = .36), condition × block (*F*(8, 322) = 1.13, *p* = .34), group × block (*F*(4, 322) = 0.95, *p* = .43), and the group × condition × block (*F*(8, 322) = 0.23, *p* = .99) interactions all failed to reach significance. The lack of interactions indicates that motor performance within a given condition was stable and did not change significantly as a function of time (across the five blocks) or group (figure 2).

**Figure 1.**
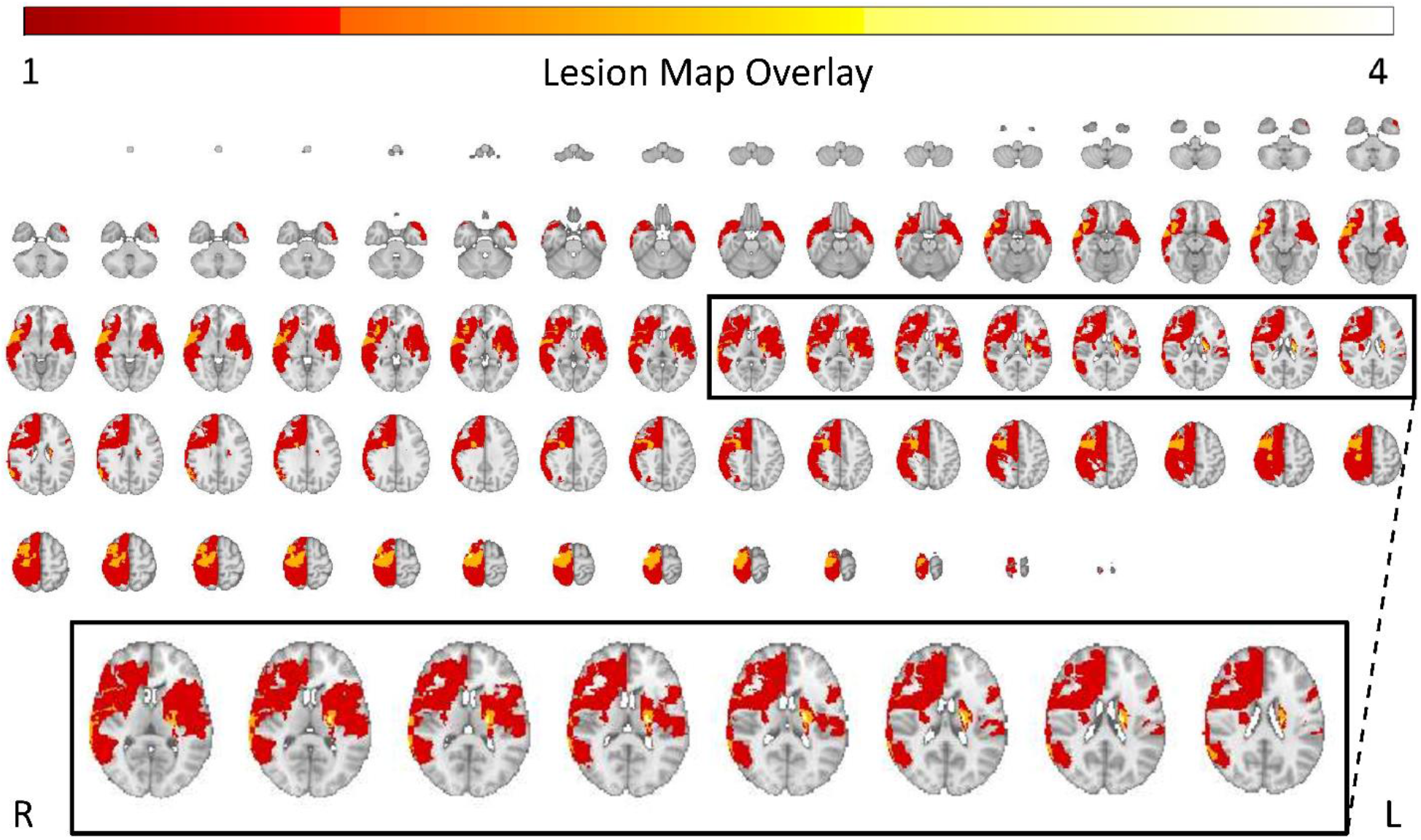
Stroke lesion map overlay. Image is in radiological view (left hemisphere displayed on the right; right hemisphere displayed on the left). Lesion masks are in MNI152 2mm standard space. Colour bar represents the number of participants that share a lesion location.

**Figure 2.**
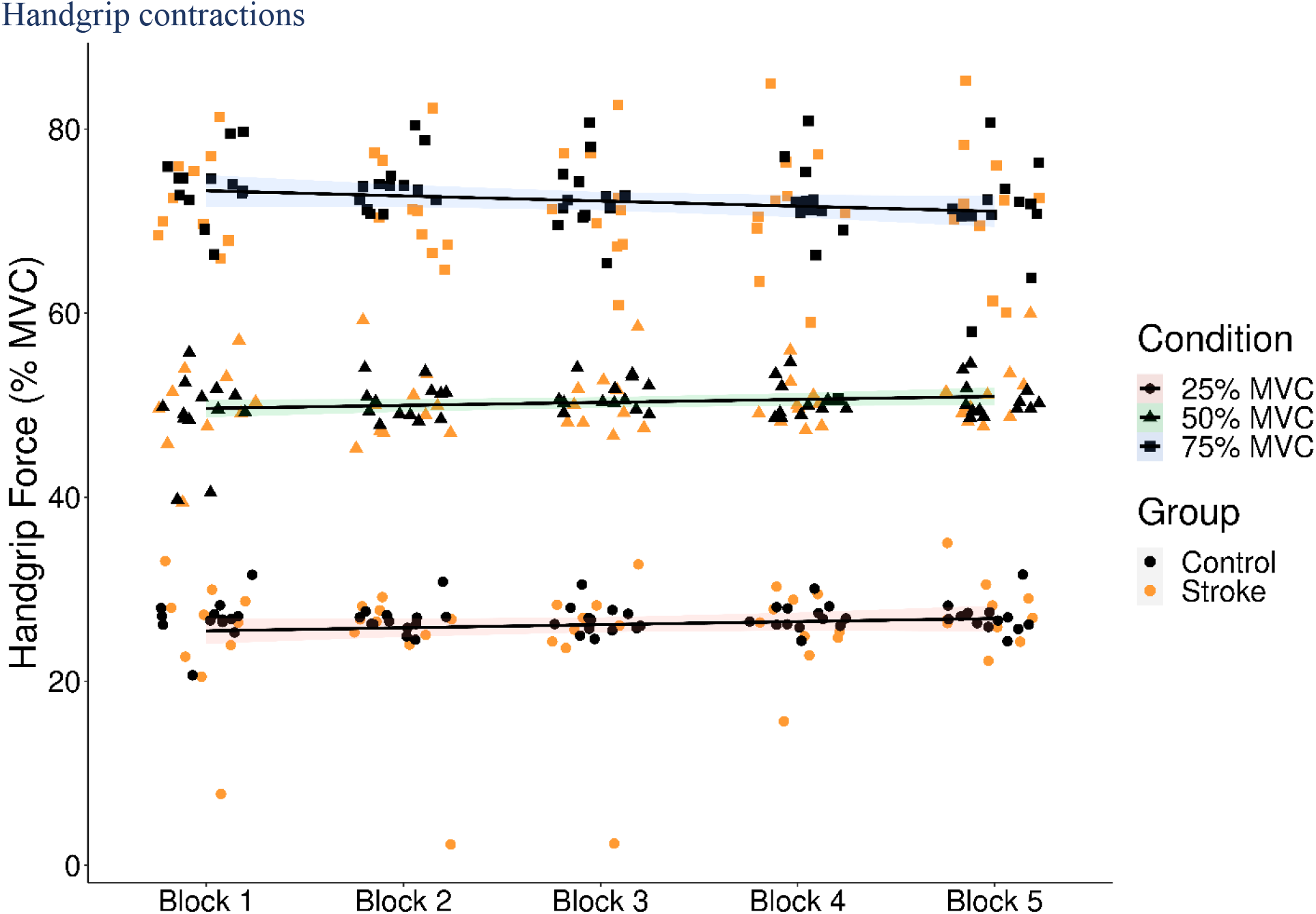
Mean relative handgrip force (y-axis: Percent MVC) for each task block (x-axis) for 25% (circles), 50% (triangles), and 75% (squares) MVC for control (black) and stroke (orange) participants. Each regression line represents each condition across blocks. Shaded area is the 95% confidence interval around the regression line.

#### Contralateral/Contralesional primary motor cortex (cM1)

For the cM1 brain activation, the main effects of condition (*F*(2, 44.69) = 2.63, *p* = .083), group (*F*(1, 23.51) = 2.28, *p* = .14) and the group × condition interaction (*F*(2, 44.69) = 0.01, *p* = .99) all did not reach significance, indicating that there were no differences in cM1 brain activation between groups or conditions (figure 3A; table 3).

**Table 3.** Group and condition means and standard deviations for the dependent variables of interest

| Variables of interest | Stroke |  |  | Healthy |  |  |
| --- | --- | --- | --- | --- | --- | --- |
|  | 25% MVC | 50% MVC | 75% MVC | 25% MVC | 50% MVC | 75% MVC |
| Force (% MVC) | 25.3 ± 6.3 | 50.3 ± 3.7 | 71.9 ± 6.1 | 26.8 ± 1.7 | 50.3 ± 2.6 | 72.4 ± 4.8 |
| iM1 (% Δ) | 0.07 ± 0.21 | 0.18 ± 0.22 | 0.26 ± 0.23 | 0.09 ± 0.25 | 0.22 ± 0.2 | 0.48 ± 0.28 |
| cM1 (% Δ) | 0.52 ± 0.36 | 0.64 ± 0.53 | 0.77 ± 0.47 | 0.69 ± 0.36 | 0.8 ± 0.24 | 0.89 ± 0.24 |
| iSMA (% Δ) | 0.35 ± 0.41 | 0.34 ± 0.38 | 0.41 ± 0.3 | 0.59 ± 0.29 | 0.69 ± 0.31 | 0.77 ± 0.28 |
| cSMA (% Δ) | 0.43 ± 0.42 | 0.48 ± 0.41 | 0.55 ± 0.36 | 0.64 ± 0.31 | 0.71 ± 0.24 | 0.77 ± 0.26 |
| cM1-iM1 (Z-Score) | 0.79 ± 0.51 | 0.85 ± 0.77 | 0.81 ± 0.46 | 0.74 ± 0.5 | 1 ± 0.27 | 1.11 ± 0.32 |
| cSMA-iSMA (Z-Score) | 1.8 ± 0.51 | 1.64 ± 0.69 | 1.54 ± 0.56 | 1.8 ± 0.42 | 2.05 ± 0.29 | 1.9 ± 0.44 |
| iM1-iSMA (Z-Score) | 1.12 ± 0.63 | 1.16 ± 0.77 | 0.93 ± 0.32 | 0.78 ± 0.51 | 1.2 ± 0.24 | 1.35 ± 0.23 |
| cM1-cSMA (Z-Score) | 1.49 ± 0.62 | 1.33 ± 0.65 | 1.38 ± 0.39 | 1.51 ± 0.43 | 1.69 ± 0.35 | 1.49 ± 0.49 |

**Figure 3.**
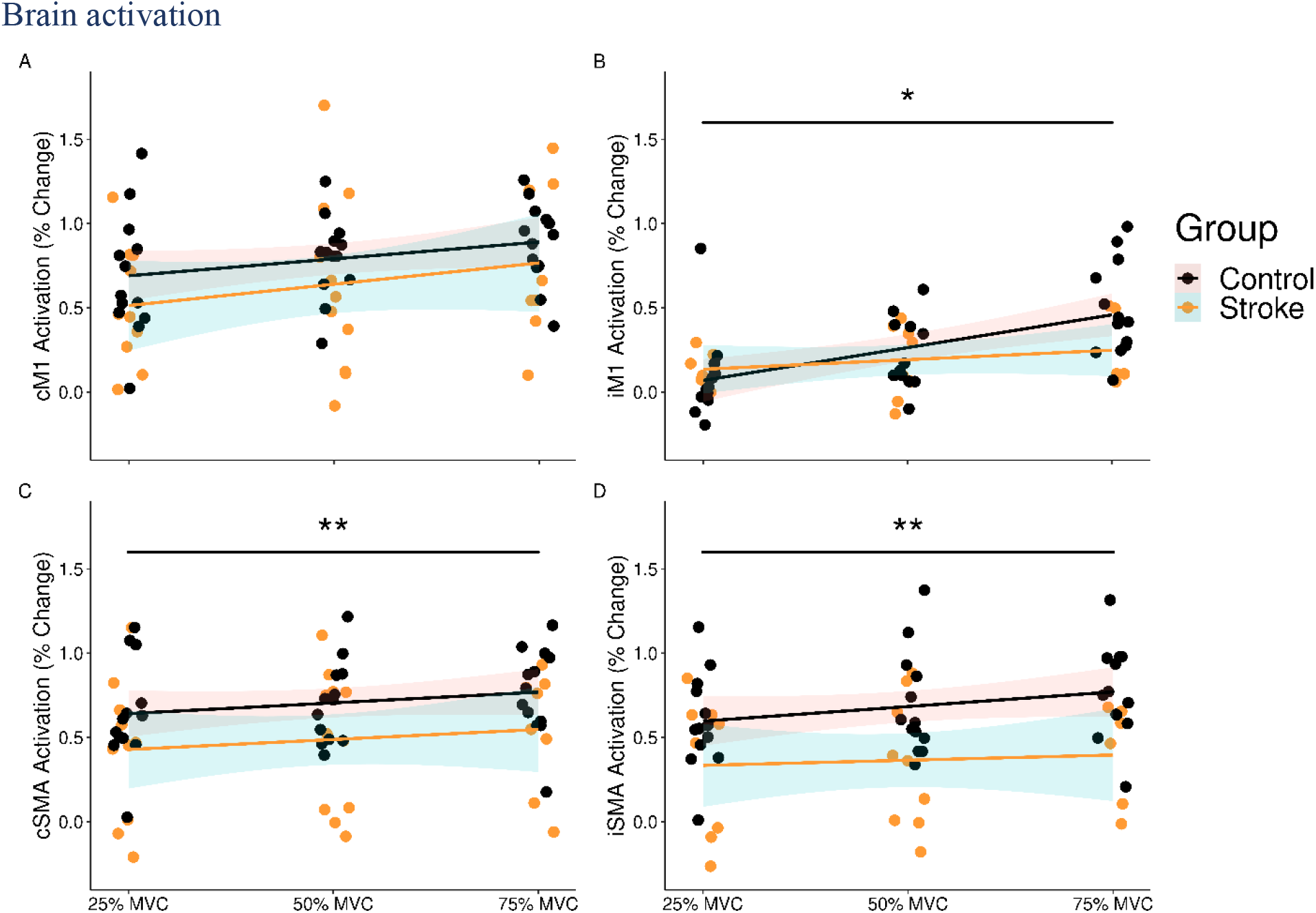
Percent change in BOLD signal for the contralateral/contralesional (c; left column), and ipsilateral/ipsilesional hemispheres (i; right column) for **A)** cM1, **B)** iM1, **C)** cSMA, **D)** iSMA for control (black) and stroke (orange) participants. * = significant main effect of condition, (*p* =.002). ** = significant main effect of group (*p* ≤ .05). Scatter plots display 95% confidence intervals around the regression line.

#### Ipsilateral/Ipsilesional primary motor cortex (iM1)

A significant main effect of condition was observed in the iM1 (*F*(2, 41.56) = 7.24, *p* = .002), where the brain activation in iM1 increased parametrically with higher force handgrip contractions. The main effect of group (*F*(1, 22.82) = 1.82, *p* = .19) and the group × condition interaction (*F*(2, 41.56) = 1.14, *p* = .33) were both non-significant, indicating that there were no differences between groups for any of the three conditions (figure 3B; table 3).

#### Contralateral/Contralesional supplementary motor area (cSMA)

For the cSMA brain activation, there was a significant main effect of group (*F*(1, 23.94) = 4.28, *p* = .050), but the main effect of condition (*F*(2, 44.87) = 1.02, *p* = .37), and the group × condition interaction (*F*(2, 44.87) = 0.09, *p* = .92) did not reach significance. These results indicate that control participants had greater cSMA brain activation compared to participants with stroke, but there were no differences between conditions for either group (figure 3C; table 3).

#### Ipsilateral/Ipsilesional supplementary motor area (iSMA)

A significant main effect of group was observed in the iSMA (*F*(1, 22.61) = 7.97, *p* = .0097), indicating that control participants had greater SMA brain activation in the ipsilateral hemisphere compared to the ipsilesional hemisphere in participants with stroke. However, the main effect of condition (*F*(2, 41.41) = 0.99, *p* = .38) and the group × condition interaction (*F*(2, 41.41) = 0.59, *p* = .56) both did not reach significance suggesting that there were no differences between conditions for either group (figure 3D; table 3).

### Functional connectivity

#### cM1-iM1 Connectivity

For cM1-iM1 functional connectivity the main effects of condition (*F*(2, 37.97) = 1.06, *p* = .36), group (*F*(1, 19.51) = 1.36, *p* = .26) and the group × condition interaction (*F*(2, 37.97) = 1.11, *p* = .34) all failed to reach significance, indicating that there were no differences between groups or conditions for cM1-iM1 interhemispheric functional connectivity (figure 4A; table 3).

**Figure 4.**
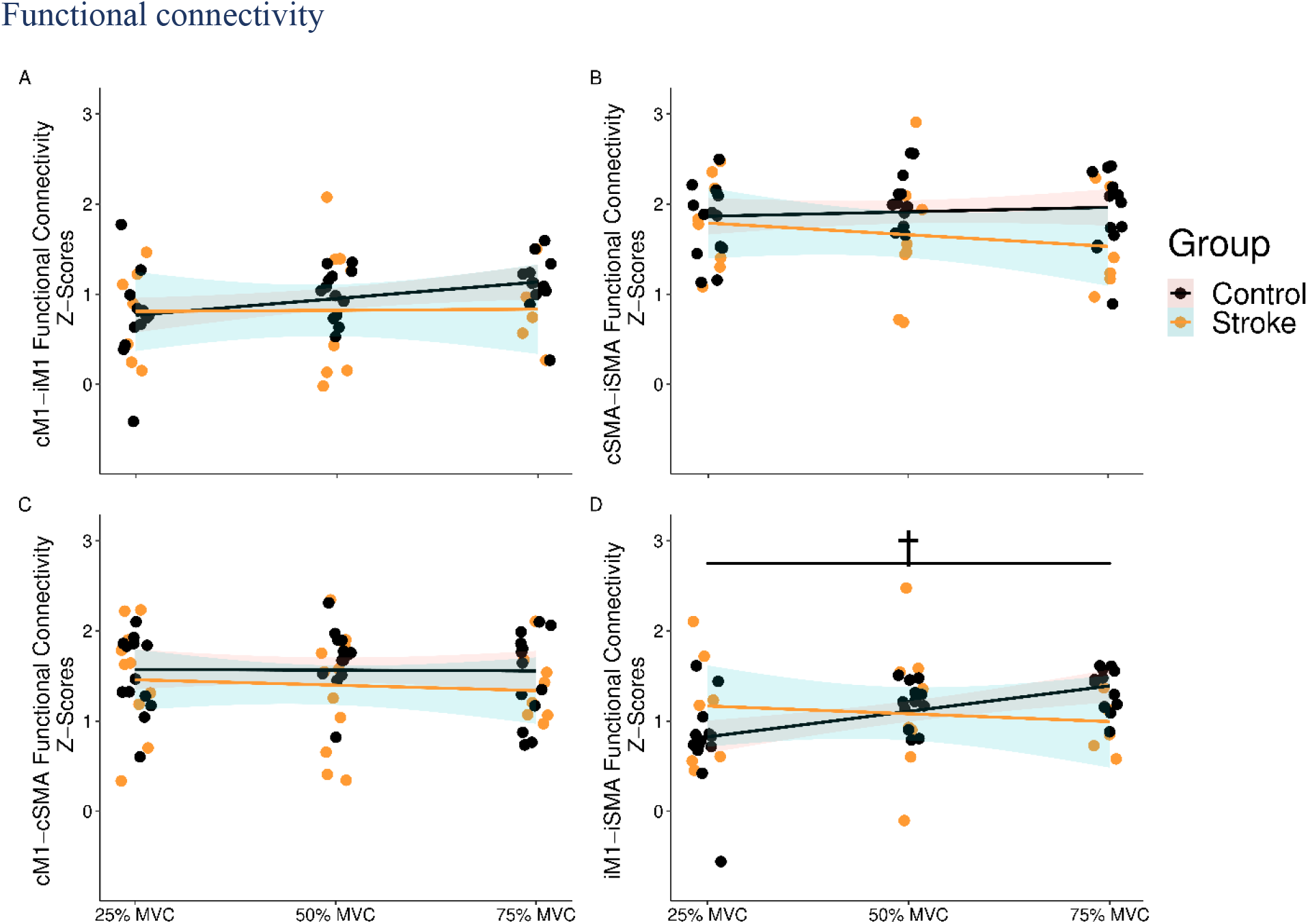
On-task functional connectivity for interhemispheric **A)** cM1-iM1, **B)** cSMA-iSMA, and intrahemispheric **C)** cM1-cSMA, and **D)** iM1-iSMA correlations. Data points are z-scores for each participant in the control (black) and stroke (orange) groups. † = significant group × condition interaction, (*p* =.027). Scatter plots display 95% confidence intervals around the regression line.

#### cSMA-iSMA Connectivity

For cSMA-iSMA functional connectivity the main effects of condition (*F*(2, 40.61) = 0.38, *p* = .69), group (*F*(1, 20.21) = 4.31, *p* = .051) and the group × condition interaction (*F*(2, 40.61) = 1.23, *p* = .30) all did not reach significance, indicating that there were no differences in cSMA-iSMA interhemispheric functional connectivity between groups or conditions (figure 4B; table 3).

#### cM1-cSMA Connectivity

For the cM1-cSMA intrahemispheric functional connectivity, the main effects of condition (*F*(2, 41.89) = 0.78, *p* = .46), and group (*F*(1, 20.73) = 1.95, *p* = .18) in addition to the group × condition interaction (*F*(2, 41.89) = 1.01, *p* = .37) all did not reach significance, indicating that this relationship did not differ regardless of group or condition (figure 4C; table 3).

#### iM1-iSMA Connectivity

The analysis of the iM1-iSMA intrahemispheric functional connectivity failed to observe significant main effects of condition (*F*(2, 37.88) = 1.58, *p* = .22), and group (*F*(1, 19.12) = 0.18, *p* = .68). However, a significant group × condition interaction was observed (*F*(2, 37.88) = 3.97, *p* = .027), indicating that iM1-iSMA functional connectivity changed differently between conditions for each group. The control group experienced a small increase in functional connectivity with increased contraction force, whereas a small decrease in connectivity was observed with increasing handgrip force for participants with stroke (figure 4D; table 3).

## Discussion

This study investigated cortical brain activation and functional connectivity during unilateral handgrip contractions with 25%, 50%, and 75% MVC. Previously published data in a neurologically intact cohort has demonstrated robust parametric scaling in the ipsilateral hemisphere with increases in handgrip force over two experimental sessions (Andrushko et al., 2021). The present study investigated this phenomenon in participants with stroke while performing the handgrip contractions with their less-affected hand and compared those findings to data from the first session of the neurologically intact control participants in previously published work (Andrushko et al., 2021). In participants with stroke, iM1 brain activation scaled with force similar to what was observed in control individuals (Andrushko et al., 2021). A notable difference between the control and participants with stroke was seen with the functional connectivity analyses. In the group of participants with stroke, inter- and intra-hemispheric functional connectivity did not scale with handgrip force as was previously observed (Andrushko et al., 2021), and with the intrahemispheric iM1-iSMA functional connectivity analyses in this study with neurologically intact control participants. This study presents novel findings that shed light on the impact of high-force unimanual contractions with the less-affected limb for promoting ipsilesional brain activity contrasting the underlying principle of the interhemispheric competition model.

There is evidence to suggest that the utilization of cross-education in participants with stroke can improve functional outcomes, whereby unilateral strength training of the less-affected limb aids in the motor recovery of the more-affected limb (Dragert and Zehr, 2013; Urbin et al., 2015; Sun et al., 2018; Dehno et al., 2021). Based on the cross-activation hypothesis for describing the neural mechanisms of cross-education, the phenomenon manifests through ipsilateral brain activation resulting in neuroplasticity that accounts for motor improvements in the contralateral limb (Lee et al., 2010; Ruddy and Carson, 2013). However, this form of motor training might be contraindicated given contralesional brain activation is thought to facilitate an inhibitory effect on the lesioned hemisphere (Kinsbourne, 1974; Murase et al., 2004; Bütefisch et al., 2008; Grefkes et al., 2008; Nowak et al., 2009). This study explicitly investigated this controversy by comparing brain activation and functional connectivity in the ipsilesional hemisphere during less-affected handgrip contractions in participants with stroke and compared those results to ipsilateral hemispheric brain activation and connectivity in neurologically intact control participants who completed right handgrip contractions.

### High-force contractions modulate the motor cortex in the ipsilateral hemisphere

The main finding from this study is that an increase in ipsilateral/ipsilesional primary motor cortex (iM1) brain activation was observed with increased handgrip force (figure 3B) in both groups. These data suggest that high-force contractions with the less-affected hand of participants with stroke did not inhibit iM1 brain activation, a possible indication that iM1 was not inhibited. Previous literature supports this notion in that the BOLD signal reflect primarily excitatory rather than inhibitory neuronal activity (Logothetis et al., 2001; Sotero and Trujillo-Barreto, 2007; Logothetis, 2008). These findings contrast the assumptions of the interhemispheric competition model, where this increased contralesional brain activation driving the less-affected handgrip contractions should have an inhibitory effect on the lesioned hemisphere. Similar to previous observations, a lack of cM1 scaling with increased contraction force is not entirely surprising (Dettmers et al., 1996; Andrushko et al., 2021). Previous literature suggests that cM1 cortical activation scales for low force contractions, but the relationship is diminished once contraction force exceeds >10% MVC (Dettmers et al., 1996; Andrushko et al., 2021). The loss of cM1 scaling with higher force contractions suggests that contraction force variance cannot be entirely accounted for with cM1 activation and corticospinal tract volleys (Cheney and Fetz, 1980; Lemon et al., 1986; Maier et al., 1993; Dettmers et al., 1996).

### Lower SMA cortical activation in participants with stroke

The SMA is considered an important cortical region for upper limb movement planning and execution (Goldberg, 1985; Forstmann et al., 2008), and is known to have direct descending connections with upper limb alpha-motoneurons (Maier et al., 2002). The SMA is also an important region in the context of stroke due to previous observations that the non-decussated reticulospinal tract serves as a compensatory pathway for motor recovery (Baker, 2011), which has cortical origins within the SMA (Fisher et al., 2021). The SMA is an important region for cross-education given its dense interhemispheric white matter connections with its homolog in the opposite hemisphere (Ruddy et al., 2017). Here we demonstrate that cortical activation in the cSMA and iSMA did not differ between conditions, suggesting this region does not modulate with force. However, a main effect of group for each region suggests that regardless of hemisphere or condition, SMA activation is lower in participants with stroke (figure 3C and D). The findings that the SMA does not modulate with force was somewhat surprising given that this region likely plays a substantial role in in stroke recovery. However, the lack of modulation does not exclude the SMA as an important cortical region for motor function. Rather, the SMA may have a more stable function in motor planning or movement generation that is not modulated by effort or force. Another possibility for the lower SMA activation and lack of modulation may be attributed to the differences in age between the participants with stroke and neurologically intact controls (stroke: mean 60 ± 11 yrs; control: 28 ± 6 yrs). Previous literature has demonstrated that with aging, atrophy occurs in the rostral segments of the corpus callosum (Hou and Pakkenberg, 2012). The SMA is located in the frontal lobe with some of the densest transcallosal fibres between homologs (Ruddy et al., 2017). Therefore, it is possible that the lower SMA activity in the participants with stroke is related to a degradation of these regions as a result of age and not due to the stroke.

### Functional connectivity is not modulated with force in participants with stroke

For the inter- and intrahemispheric functional connectivity analyses, the only significant observation was a group × condition interaction for the iM1-iSMA connectivity (*p* = .027). This interaction was influenced by an increase in connectivity strength with higher force contractions in the control participants only, whereas no change was observed in the participants with stroke (figure 4D). The only other finding that was nearly significant was the main effect of group for cSMA-iSMA functional connectivity (*p* = .051), whereby control participants displayed marginally higher levels of functional connectivity than participants with stroke across the three conditions (table 3). Overall, these data suggest that inter- and intrahemispheric functional connectivity are not significantly modulated with contraction force in participants with stroke. It remains to be determined if the lack of modulation paired with the lower connectivity across conditions for participants with stroke are related to a degradation in motor function, or whether the observed difference in functional connectivity is simply related to aging where differences have been observed (Tscherpel et al., 2020).

#### Limitations and future directions

These data provide an important foundation for future work in understanding neural correlates of unimanual motor tasks with the less-affected limb in participants with stroke, but there are several important limitations in this study. First, because the study used neurologically intact controls from a previous study (Andrushko et al., 2021), they were not aged-matched with the stroke group, perhaps limiting comparisons between groups. Future research should aim to replicate these findings with age-matched neurologically intact controls. Measuring brain activity during high-force contractions is difficult with fMRI, given that head motion can contaminate the data and render it unusable (Makowski et al., 2019). Further, high intensity or high-force strength-based exercises are not conventionally used in stroke rehabilitation, with most rehabilitation programs focused primarily on motor skill recovery (Langhorne et al., 2009; Belagaje, 2017). Given these two points, current literature is lacking on the fundamental understanding of how high-force, strength-based exercise may modulate neuroplasticity and promote motor recovery in stroke. With that in mind, an additional limitation was the relatively low sample size within each group. Recruiting participants with stroke to participate in research is difficult and future work may benefit from collaborative and multi-site studies to increase the sample size to offset data loss with these high-force motor tasks in the fMRI environment. Further, an inspection of the lesion volume (table 1) and location (figure 1) clearly shows that study participants were a heterogenous sample of participants with stroke, with a maximum lesion overlap of four (i.e., number of participants with a shared lesion location). Future multi-site collaborative efforts may be able to better screen participants to obtain a more homogenous group in terms of lesion size, location, time since stroke, and functional scores, which was limited in the present study in part due to the impact of the global COVID-19 pandemic on participant recruitment. An additional limitation is that performing high-force handgrip contractions during fMRI brain scans is difficult to achieve without substantial data loss due to motion contamination. We contend that the cost-benefit trade-off to carrying out investigations into how the brain functions during these higher force contractions are of value and imperative for scientific and clinical advancement.

## Conclusion

This study suggests high-force contractions with the less-affected limb may provide greater benefit to promoting ipsilateral/ipsilesional (iM1) activity subserving use-dependent neuroplasticity than lower force contractions. Further, the modulation of iM1 brain activation does not appear to be influenced by interhemispheric communication between these homologous regions or intrahemispheric connectivity between the SMA and M1, and is likely influenced independently or by other inter- or intrahemispheric connections not measured in the present study. The study provides a possible mechanistic basis for which cross-education may be utilized to promote iM1 brain activity in participants with stroke. In scenarios where individuals with stroke do not have the functional capacity to engage in constraint-induced-movement-therapy, whereby the more-affected limb is directly exercised (Grotta et al., 2004), unilateral high-force contractions with the less-affected limb can provide a boost in iM1 activity that may lead to neuroplasticity of the lesioned M1 circuitry.

## Acknowledgements

We would like to acknowledge the excellent work of Sharleen Weese Maley, Research Program Manager, Clinical Trial Support Unit and Saskatchewan Cerebrovascular Centre. We thank Mr. Shawn Reinink from the University of Saskatchewan for his assistance in LabView programming, Jake Mota, Ph.D. from the University of Alabama, and Caroline Nettekoven, Ph.D. from the University of Cambridge for their guidance in carrying out Linear Mixed Effects Analyses in R.

## References

Andersson JLR, Jenkinson M, Smith S (2007a) Non-linear registration aka spatial normalisation FMRIB Technial Report TR07JA2.

Andersson JLR, Jenkinson M, Smith S (2007b) Non-linear optimisation FMRIB Technial Report TR07JA1.

Andrushko JW, Gould LA, Renshaw DW, Ekstrand C, Hortobágyi T, Borowsky R, Farthing JP (2021) High force unimanual handgrip contractions increase ipsilateral sensorimotor activation and functional connectivity. Neuroscience 452:111–125.

Baker SN (2011) The primate reticulospinal tract, hand function and functional recovery. J Physiol 589:5603–5612 Available at: http://www.ncbi.nlm.nih.gov/pubmed/21878519 [Accessed September 5, 2019].

Belagaje SR (2017) Stroke rehabilitation. Contin Lifelong Learn Neurol 23:238–253 Available at: https://journals.lww.com/continuum/Fulltext/2017/02000/Stroke_Rehabilitation.17.aspx [Accessed May 15, 2021].

Benwell NM, Sacco P, Hammond GR, Byrnes ML, Mastaglia FL, Thickbroom GW (2006) Short-interval cortical inhibition and corticomotor excitability with fatiguing hand exercise: a central adaptation to fatigue? Exp Brain Res 170:191–198 Available at: http://link.springer.com/10.1007/s00221-005-0195-7.

Borowsky R, Owen WJ, Sarty GE (2002) The role of the left hemisphere in motor control of touch: A functional magnetic resonance imaging analysis. Brain Cogn 49:96–101 Available at: https://pubmed.ncbi.nlm.nih.gov/12027395/ [Accessed January 4, 2021].

Bryden MP (1977) Measuring handedness with questionnaires. Neuropsychologia 15:617–624.

Buetefisch CM (2015) Role of the contralesional hemisphere in post-stroke recovery of upper extremity motor function. Front Neurol 6:214 Available at: http://www.ncbi.nlm.nih.gov/pubmed/26528236 [Accessed September 3, 2019].

Bütefisch CM, Weßling M, Netz J, Seitz RJ, Hömberg V (2008) Relationship between interhemispheric inhibition and motor cortex excitability in subacute stroke patients. Neurorehabil Neural Repair 22:4–21 Available at: https://journals.sagepub.com/doi/abs/10.1177/1545968307301769 [Accessed May 23, 2021].

Cheney PD, Fetz EE (1980) Functional classes of primate corticomotoneuronal cells and their relation to active force. J Neurophysiol 44:773–791 Available at: https://journals.physiology.org/doi/abs/10.1152/jn.1980.44.4.773 [Accessed May 23, 2021].

Cincotta M, Ziemann U (2008) Neurophysiology of unimanual motor control and mirror movements. Clin Neurophysiol 119:744–762.

Cole MW, Ito T, Schultz D, Mill R, Chen R, Cocuzza C (2018) Task activations produce spurious but systematic inflation of task functional connectivity estimates. Neuroimage 189:1–18 Available at: http://www.ncbi.nlm.nih.gov/pubmed/30597260.

Dehno SN, Kamali F, Shariat A, Jaberzadeh S (2021) Unilateral strength training of the less affected hand improves cortical excitability and clinical outcomes in patients with subacute stroke: A randomized controlled trial. Arch Phys Med Rehabil 102 Available at: https://pubmed.ncbi.nlm.nih.gov/33460575/ [Accessed April 28, 2021].

Dettmers C, Ridding MC, Stephan KM, Lemon RN, Rothwell JC, Frackowiak RSJ (1996) Comparison of regional cerebral blood flow with transcranial magnetic stimulation at different forces. J Appl Physiol 81:596–603 Available at: https://pubmed.ncbi.nlm.nih.gov/8872623/ [Accessed May 23, 2021].

Dodd KC, Nair VA, Prabhakaran V (2017) Role of the contralesional vs. ipsilesional hemisphere in stroke recovery. Front Hum Neurosci 11:469 Available at: http://www.ncbi.nlm.nih.gov/pubmed/28983244 [Accessed September 18, 2019].

Dragert K, Zehr EP (2013) High-intensity unilateral dorsiflexor resistance training results in bilateral neuromuscular plasticity after stroke. Exp Brain Res 225:93–104.

Fan L, Li H, Zhuo J, Zhang Y, Wang J, Chen L, Yang Z, Chu C, Xie S, Laird AR, Fox PT, Eickhoff SB, Yu C, Jiang T (2016) The human brainnetome atlas: A new brain atlas based on connectional architecture. Cereb Cortex 26:3508–3526 Available at: http://www.ncbi.nlm.nih.gov/pubmed/27230218 [Accessed April 17, 2020].

Farthing JP, Chilibeck PD (2003) The effect of eccentric training at different velocities on cross-education. Eur J Appl Physiol 89:570–577.

Fisher KM, Zaaimi B, Edgley SA, Baker SN (2021) Extensive cortical convergence to primate reticulospinal pathways. J Neurosci 41:1005–1018 Available at: https://pubmed.ncbi.nlm.nih.gov/33268548/ [Accessed May 23, 2021].

Forstmann BU, Dutilh G, Brown S, Neumann J, Von Cramon DY, Ridderinkhof KR, Wagenmakers EJ (2008) Striatum and pre-SMA facilitate decision-making under time pressure. Proc Natl Acad Sci U S A 105:17538–17542 Available at: www.pnas.org/cgi/content/full/ [Accessed January 24, 2021].

Goldberg G (1985) Supplementary motor area structure and function: Review and hypotheses. Behav Brain Sci 8:567–588.

Gowland C, Stratford P, Ward M, Moreland J, Torresin W, Van Hullenaar S, Sanford J, Barreca S, Vanspall B, Plews N (1993) Measuring physical impairment and disability with the chedoke-mcmaster stroke assessment. Stroke 24:58–63 Available at: http://ahajournals.org [Accessed May 18, 2021].

Grefkes C, Eickhoff SB, Nowak DA, Dafotakis M, Fink GR (2008) Dynamic intra- and interhemispheric interactions during unilateral and bilateral hand movements assessed with fMRI and DCM. Neuroimage 41.

Grotta JC, Noser EA, Ro T, Boake C, Levin H, Aronowski J, Schallert T (2004) Constraint-induced movement therapy. In: Stroke, pp 2699–2701. Lippincott Williams & Wilkins. Available at: http://stroke.ahajournals.org/cgi/doi/10.1161/01.STR.0000143320.64953.c4 [Accessed January 2, 2021].

Hou J, Pakkenberg B (2012) Age-related degeneration of corpus callosum in the 90+ years measured with stereology. Neurobiol Aging 33:1009.e1–1009.e9 Available at: https://pubmed.ncbi.nlm.nih.gov/22118947/ [Accessed June 8, 2021].

Hübers A, Orekhov Y, Ziemann U (2008) Interhemispheric motor inhibition: Its role in controlling electromyographic mirror activity. Eur J Neurosci 28:364–371 Available at: https://onlinelibrary.wiley.com/doi/full/10.1111/j.1460-9568.2008.06335.x [Accessed May 21, 2021].

Jenkinson M, Bannister P, Brady M, Smith S (2002) Improved optimization for the robust and accurate linear registration and motion correction of brain images. Neuroimage 17:825–841 Available at: http://www.ncbi.nlm.nih.gov/pubmed/12377157.

Jenkinson M, Smith S (2001) A global optimisation method for robust affine registration of brain images. Med Image Anal 5:143–156 Available at: http://www.ncbi.nlm.nih.gov/pubmed/11516708.

Jiang Z, Wang X-F, Kisiel-Sajewicz K, Yan JH, Yue GH (2012) Strengthened functional connectivity in the brain during muscle fatigue. Neuroimage 60:728–737 Available at: http://linkinghub.elsevier.com/retrieve/pii/S1053811911014157.

Katan M, Luft A (2018) Global burden of stroke. Semin Neurol 38:208–211 Available at: www.zora.uzh.chyear:2018URL:https://doi.org/10.5167/uzh [Accessed May 3, 2021].

Kinsbourne M (1974) Mechanisms of hemispheric interaction in man. Hemispheric Disconnection Cereb Funct XIII.

Kuznetsova A, Brockhoff PB, Christensen RHB (2017) lmerTest Package: Tests in Linear Mixed Effects Models . J Stat Softw 82.

Langhorne P, Coupar F, Pollock A (2009) Motor recovery after stroke: a systematic review. Lancet Neurol 8:741–754.

Lee M, Carroll TJ (2007) Cross education: Possible mechanisms for the contralateral effects of unilateral resistance training. Sport Med 37:1–14.

Lee M, Hinder MR, Gandevia SC, Carroll TJ (2010) The ipsilateral motor cortex contributes to cross-limb transfer of performance gains after ballistic motor practice. J Physiol 588:201–212.

Lemon RN, Mantel GW, Muir RB (1986) Corticospinal facilitation of hand muscles during voluntary movement in the conscious monkey. J Physiol 381:497–527 Available at: https://pubmed.ncbi.nlm.nih.gov/3625543/ [Accessed May 23, 2021].

Logothetis NK (2008) What we can do and what we cannot do with fMRI. Nature 453:869–878 Available at: https://pubmed.ncbi.nlm.nih.gov/18548064/ [Accessed June 16, 2021].

Logothetis NK, Pauls J, Augath M, Trinath T, Oeltermann A (2001) Neurophysiological investigation of the basis of the fMRI signal. Nature 412:150–157 Available at: https://pubmed.ncbi.nlm.nih.gov/11449264/ [Accessed June 16, 2021].

Maier MA, Armand J, Kirkwood PA, Yang H-W, Davis JN, Lemon R.N. (2002) Differences in the corticospinal projection from primary motor cortex and supplementary motor area to macaque upper limb motoneurons: An anatomical and electrophysiological study. Cereb Cortex 12:281–296 Available at: https://academic.oup.com/cercor/article-lookup/doi/10.1093/cercor/12.3.281 [Accessed December 26, 2020].

Maier MA, Bennett KMB, Hepp-Reymond MC, Lemon RN (1993) Contribution of the monkey corticomotoneuronal system to the control of force in precision grip. J Neurophysiol 69:772–785 Available at: https://pubmed.ncbi.nlm.nih.gov/8463818/ [Accessed May 23, 2021].

Makowski C, Lepage M, Evans AC (2019) Head motion: The dirty little secret of neuroimaging in psychiatry. J Psychiatry Neurosci 44:62–68 Available at: /pmc/articles/PMC6306289/ [Accessed May 15, 2021].

Manca A, Dragone D, Dvir Z, Deriu F (2017) Cross-education of muscular strength following unilateral resistance training: a meta-analysis. Eur J Appl Physiol 117:2335–2354 Available at: http://link.springer.com/10.1007/s00421-017-3720-z [Accessed January 29, 2018].

Manca A, Hortobágyi T, Carroll TJ, Enoka RM, Farthing JP, Gandevia SC, Kidgell DJ, Taylor JL, Deriu F (2021) Contralateral effects of unilateral strength and skill training: Modified delphi consensus to establish key aspects of cross-education. Sport Med 51:11–20 Available at: https://doi.org/10.1007/s40279-020-01377-7 [Accessed January 23, 2021].

Manca A, Hortobágyi T, Rothwell J, Deriu F (2018) Neurophysiological adaptations in the untrained side in conjunction with cross-education of muscle strength: a systematic review and meta-analysis. J Appl Physiol 124:1502–1518 Available at: https://www.physiology.org/doi/10.1152/japplphysiol.01016.2017.

Mittmann N, Seung SJ, Hill MD, Phillips SJ, Hachinski V, Coté R, Buck BH, Mackey A, Gladstone DJ, Howse DC, Shuaib A, Sharma M (2012) Impact of disability status on ischemic stroke costs in Canada in the first year. Can J Neurol Sci / J Can des Sci Neurol 39:793–800 Available at: https://www.cambridge.org/core/product/identifier/S0317167100015638/type/journal_article [Accessed September 3, 2019].

Murase N, Duque J, Mazzocchio R, Cohen LG (2004) Influence of interhemispheric interactions on motor function in chronic stroke. Ann Neurol 55:400–409 Available at: https://pubmed.ncbi.nlm.nih.gov/14991818/ [Accessed May 23, 2021].

Nowak DA, Grefkes C, Ameli M, Fink GR (2009) Interhemispheric competition after stroke: Brain stimulation to enhance recovery of function of the affected hand. Neurorehabil Neural Repair 23:641–656 Available at: http://www.ncbi.nlm.nih.gov/pubmed/19531606 [Accessed September 3, 2019].

Pruim RHR, Mennes M, van Rooij D, Llera A, Buitelaar JK, Beckmann CF (2015) ICA-AROMA: A robust ICA-based strategy for removing motion artifacts from fMRI data. Neuroimage 112:267–277.

R Core Team (2019) R: A language and environment for statistical computing (Version 3.6). R Found Stat Comput.

Ruddy KL, Carson RG (2013) Neural pathways mediating cross education of motor function. Front Hum Neurosci 7:1–22 Available at: http://journal.frontiersin.org/article/10.3389/fnhum.2013.00397/abstract.

Ruddy KL, Leemans A, Carson RG (2017) Transcallosal connectivity of the human cortical motor network. Brain Struct Funct 222:1243–1252.

Scripture EW, Smith TL, Brown EM (1894) On the education of muscular control and power. Stud from Yale Psychol Lab:114–119 Available at: http://echo.mpiwg-berlin.mpg.de/ECHOdocuView?url=/permanent/vlp/lit23174/index.meta [Accessed January 29, 2018].

Sehm B, Steele CJ, Villringer A, Ragert P (2016) Mirror motor activity during right-hand contractions and its relation to white matter in the posterior midbody of the corpus callosum. Cereb Cortex 26:4347–4355 Available at: https://pubmed.ncbi.nlm.nih.gov/26400922/ [Accessed May 19, 2021].

Sleegers WWA (2020) tidystats: Save output of statistical tests (Version 0.5).

Smith SM (2002) Fast robust automated brain extraction. Hum Brain Mapp 17:143–155 Available at: http://www.ncbi.nlm.nih.gov/pubmed/12391568.

Sotero RC, Trujillo-Barreto NJ (2007) Modelling the role of excitatory and inhibitory neuronal activity in the generation of the BOLD signal. Neuroimage 35:149–165.

Steel A, Song S, Bageac D, Knutson KM, Keisler A, Saad ZS, Gotts SJ, Wassermann EM, Wilkinson L (2016) Shifts in connectivity during procedural learning after motor cortex stimulation: A combined transcranial magnetic stimulation/functional magnetic resonance imaging study. Cortex 74:134–148 Available at: http://www.ncbi.nlm.nih.gov/pubmed/26673946.

Sullivan KJ, Tilson JK, Cen SY, Rose DK, Hershberg J, Correa A, Gallichio J, McLeod M, Moore C, Wu SS, Duncan PW (2011) Fugl-meyer assessment of sensorimotor function after stroke: Standardized training procedure for clinical practice and clinical trials. Stroke 42:427–432 Available at: http://stroke.ahajournals.org/cgi/content/full/STROKEAHA.110.592766/DC1. [Accessed May 18, 2021].

Sun Y, Ledwell NMH, Boyd LA, Zehr EP (2018) Unilateral wrist extension training after stroke improves strength and neural plasticity in both arms. Exp Brain Res 236:2009–2021 Available at: http://link.springer.com/10.1007/s00221-018-5275-6.

The Mathworks Inc (2018) MATLAB 2018a. [WwwMathworksCom/Products/Matlab2](https://Www.Mathworks.Com/Products/Matlab2).

Tscherpel C, Hensel L, Lemberg K, Freytag J, Michely J, Volz LJ, Fink GR, Grefkes C (2020) Age affects the contribution of ipsilateral brain regions to movement kinematics. Hum Brain Mapp 41:640–655 Available at: http://www.neurobs.com [Accessed June 3, 2021].

Urbin MA, Harris-Love ML, Carter AR, Lang CE (2015) High-intensity, unilateral resistance training of a non-paretic muscle group increases active range of motion in a severely paretic upper extremity muscle group after stroke. Front Neurol 6:119 Available at: http://www.frontiersin.org/Stroke/10.3389/fneur.2015.00119/abstract.

Verstynen T, Diedrichsen J, Albert N, Aparicio P, Ivry RB (2005) Ipsilateral motor cortex activity during unimanual hand movements relates to task complexity. J Neurophysiol 93:1209–1222.

Wickham H (2016) ggplot2: Elegant graphics for data analysis.

Woolrich MW, Ripley BD, Brady M, Smith SM (2001) Temporal autocorrelation in univariate linear modeling of FMRI data. Neuroimage 14:1370–1386 Available at: http://www.ncbi.nlm.nih.gov/pubmed/11707093.

Worsley K (2001) Statistical analysis of activation images. In: Functional MRI: An Introduction to Methods, First edit. (Jezzard P, Matthews PM, Smith SM, eds), pp 251–270. Oxford: Oxford University Press.

Yu Z, Prado R, Quinlan EB, Cramer SC, Ombao H (2016) Understanding the impact of stroke on brain motor function: A hierarchical bayesian approach. J Am Stat Assoc 111:549–563 Available at: http://www.ncbi.nlm.nih.gov/pubmed/28138206 [Accessed October 2, 2019].

